# Localized Recombination Drives Diversification of Killing Spectra for Phage Derived Syringacins

**DOI:** 10.1101/240762

**Authors:** David A. Baltrus, Meara Clark, Caitlin Smith, Kevin L. Hockett

**Affiliations:** School of Plant Sciences, University of Arizona, Tucson AZ 85721; School of Animal and Comparative Biomedical Sciences, University of Arizona, Tucson AZ 85721

## Abstract

To better understand the potential for detrimental interactions between strains of the same bacterial species, we have surveyed bacteriocin killing activity across a diverse suite of strains of the phytopathogen *Pseudomonas syringae.* Our data demonstrate that killing activity from phage derived bacteriocins of *P. syringae* (R-type syringacins) is widespread. Despite a high overall diversity of bacteriocin activity, strains can broadly be classified into five main killing types and two main sensitivity types. Furthermore, we show that killing activity switches frequently between strains, and that switches correlate with localized recombination of two genes that together encode the proteins that specify bacteriocin targeting. Lastly, we demonstrate that phage derived bacteriocin killing activity can be swapped between strains simply through expression of these two genes *in trans.* Overall, our study characterizes extensive diversity of killing activity for phage derived bacteriocins of *P. syringae* across strains and highlights the power of localized recombination to alter phenotypes that mediate strain interactions during evolution of natural populations and communities.

## Introduction

Microbiome composition is dynamic through both space and time, but many questions remain concerning how specific factors shape these communities especially in the context of phytobiomes. Interactions mediated by the production of and sensitivity to molecules made by members of the community are known to play an important role in structuring microbiomes (Coyte et al., 2015; Koskella et al., 2017). Given the potential for rapid evolution of pathways underlying these interactions, they may also be particular hotbeds for coevolution within microbiomes (Bucci et al., 2011; Hawlena et al., 2012; Riley and Wertz, 2002). Furthermore, heightened appreciation for the importance of microbes in driving natural biogeochemical cycles, with greater understanding of microbial influence in host physiology and development, and with a burgeoning role for microbes across industrial systems has driven increased interest in the development of ways to exploit these factors to precisely engineer microbiomes (Mueller and Sachs, 2015; Sheth et al., 2016). A more thorough understanding of underlying patterns and of the genetic basis of interactions between microbes will therefore shed new light into mechanisms that structure ecological and evolutionary dynamics across natural microbial communities and may enable development of targeted therapies to precisely manipulate microbial composition across hosts and environments.

The best characterized, and perhaps best known, means to alter microbial community composition involves the use of broad spectrum antibiotics targeting physiological processes conserved across diverse bacterial taxa. However, lowered efficacy of broad spectrum antibiotics due to the evolution of resistance coupled with the realization that their use can lead to detrimental off-target effects on beneficial microbes has created new research momentum to identify and characterize new types of intermicrobial interactions with higher specificity (Andersson and Hughes, 2014; Cotter et al., 2013; Gebhart et al., 2015; Yang et al., 2014). Bacteriocins are largely thought to be molecules produced by bacterial cells that specifically target different strains of the same species or closely related species (Cotter et al., 2013; Riley and Wertz, 2002). In contrast to relatively indiscriminate activity of some broad spectrum antibiotics, bacteriocin targeting is mediated through a requirement for interactions with particular receptors on target cells before antimicrobial functions (Riley and Wertz, 2002; Williams et al., 2008). The precision of bacteriocin killing activity also suggests that they are used as a means to outcompete microbes that occupy similar niches, and thus characterization of bacteriocin activity and sensitivity from a diverse collection of strains could shed light on previously unrecognized ecological patterns (Hert et al., 2005). Perhaps most relevant for their therapeutic use, the relatively specific killing spectrum of bacteriocins could provide a powerful means to precisely engineer microbiomes while avoiding off-target effects seen in other treatments (Cotter et al., 2013; Sheth et al., 2016).

We have recently characterized a class of phage derived bacteriocins produced by the plant pathogen *Pseudomonas syringae,* termed R-type syringacins (Hockett et al., 2015). These compounds are a member of a broader class of molecules, called tailocins, the production of which is controlled by bacteria but which are evolutionarily derived from coopted prophage (Ghequire and De Mot, 2015). While phage derived syringacins appear to be mechanistically and evolutionarily similar to R-type pyocins produced by *P. aeruginosa,* and are indeed found at the same genomic context (in between genes involved in tryptophan biosynthesis), they have a different evolutionary history as they were derived from a different progenitor prophage (Hockett et al., 2015). Although bacteriocin activity has been characterized across subsets of *P. syringae* strains, and while it is likely that much of the reported activity was due to phage derived syringacins, earlier experiments were limited in their ability to analyze the genetic basis of changes in killing spectra because genome sequences were not available (Vidaver, 1976; Vidaver and Buckner, 1978). The relative ease of modern genome sequencing has enabled a more thorough understanding of genomic evolution and phylogenetic relationships between strains of *P.syringae (Baltrus et al., 2017).* We therefore sought to take advantage of this wealth of genomic data to characterize bacteriocin killing spectra and sensitivity across a diverse range of *P. syringae* strains in order to gain insights into evolutionary patterns and the genetic basis of phenotypic diversity in bacteriocin activity across this species.

Here we report the characterization of killing and sensitivity spectra of bacteriocins across a diverse panel of strains classified as *P. syringae* sensu lato. We find that many of the strains can be broadly grouped into two main killing activity and sensitivity classes, and that membership across these groups is correlated. We further investigate the genetic basis of switches in killing activity for multiple strain pairs, and find that such switches are likely due to localized recombination of a handful of genes in the R-type syringacin locus. Lastly, we confirm that ectopic expression of just two genes is sufficient to switch phage derived bacteriocin targeting in at least one pair of closely related strains. Our results highlight the important role that recombination plays in shaping a trait that likely influences community composition for this cosmopolitan environmental bacterium and paves the way for future studies to investigate the ecological and evolutionary basis of bacteriocin production and sensitivity across *P. syringae.*

## Results

### Phylogenetic biases in syringacin production and killing

We have performed an all all by all screen across a diverse array of *P. syringae* strains to investigate production of and sensitivity to antimicrobial compounds induced by DNA damaging agents. We have further used polyethylene glycol (PEG) precipitation to select for the subset of these compounds likely to be phage tail derived bacteriocins known as R-type syringacins. Upon inspection of the killing matrix (Figure 1) it is apparent that, although many of these strains harbor a variety of bacteriocins besides R-type syringacins, the dominant killing activity across all strains precipitates with PEG and thus is likely a high molecular weight compound. Furthermore, we note that killing activity of this high molecular weight compound was usually associated with crisp borders in agar overlay assays, which is strongly suggestive that these compounds are truly R-type syringacins (data not shown). This result echoes a screen of bacteriocin killing capability performed on a more limited collection of strains, and points to an important role for R-type syrigacins in structuring communities of *P. syringae (Hockett et al., 2015).*

**Figure 1.**
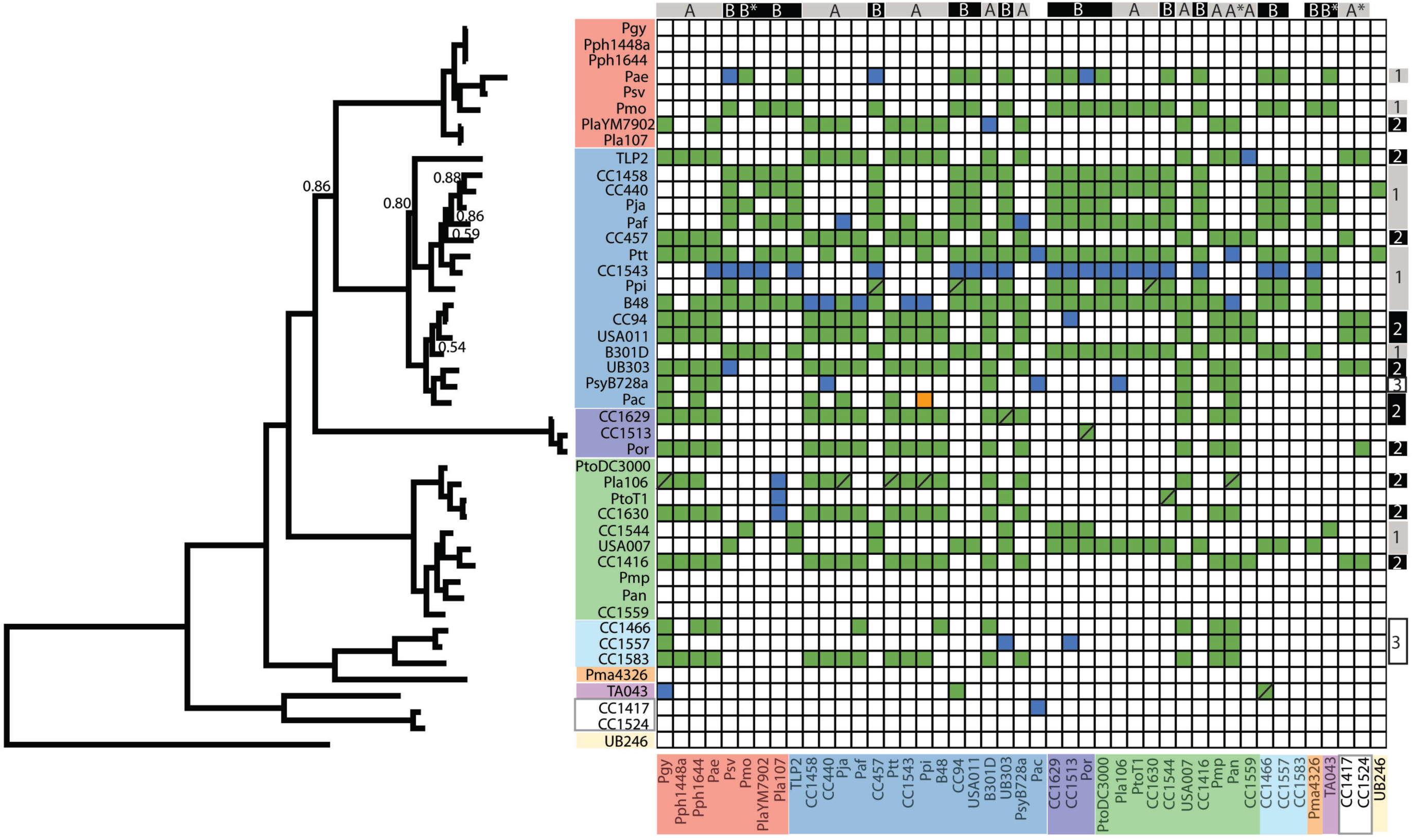
Extensive R-type syringacin killing activity across diverse *P. syringae* strains. Strains are listed in both axes according to phylogenetic relationships. On the left of the figure is a phylogeny inferred from concatenated fragments of nucleotide sequences for five genes typically used in MLSA analysis of *P. syringae (gyrB, rpoD, gapA, pgi, acnB).* Posterior probabilities <0.95 are shown. Strain names are colored on both axes by their traditional placement into phylogroups. The X-axis of the matrix displays strain sensitivities to killings, while the Y-axis displays strain killing activity. Colored boxes indicate killing activity according to overlay assays. Solid boxes indicate repeatable killing activity across multiple assays, whereas boxes with slashes indicate that killing activity existed in at least one assay. Blue boxes indicate killing activity only present in pure supernatants and thus consistent with diffusible bacteriocins, while green boxes indicate killing activity that was present in PEG selected supernatants and is thus consistent with phage derived bacteriocins. The orange box indicates killing activity that was repeatedly witnessed only after PEG precipitation. On the top of the matrix, strains have been grouped into sensitivity clusters, while on the right side of the matrix strains are grouped into killing clusters.

We have ordered the killing matrix by phylogeny, and colored strains according to phylogroup as described in (Berge et al., 2014)), so that proximity of producer and target strains along with coloration provides a readout on overall evolutionary similarity (Figure 1). Closer inspection of the overall killing matrix in the context of phylogenetic relationships reveals numerous clear patterns as well as a handful of important yet nuanced results. A far majority of strains exhibits killing activity consistent with phage derived bacteriocins to at least one other strain in this screened collection (31/45, 68%). In particular, strains within phylogroup II (14 of 15 strains) seems to be exceptionally capable of killing other *P. syringae* strains within our collection, as almost all of the tested members can target at least one other screened strain. In no case did we find that strains demonstrated killing activity against themselves. Furthermore, although they are the minority, multiple strains show no killing activity against our strain collection and these strains generally clustered by phylogeny. For instance, two strains sampled from phylogroup IX (CC1417 and CC1524) do not show any killing activity consistent with R-type syringacins. Likewise, multiple closely related strains from phylogroup I *(Pmp, Pan,* CC1559) as well as phylogroup III *(Pgy,* Pph1448a, Pph1644) display no killing activity. In each of these cases, strains that lack killing activity form monophyletic groups compared to other strains within our screened collection.

### R-type syringacin killing activity can be broadly grouped into two high activity clusters and at least five total groups

As an additional step to characterize killing spectra, we used hierarchical clustering to group strains based off of likely R-type syringacin killing activity (Figure 2). According to groupings arising from this analysis, screened strains can be sorted into at least two deeply branching main clusters (which we term clusters I and II) and at least five total groups. The two main clusters are dominated by strains with relatively broad killing spectrum (an average of ~19 and ~16 strains targeted per cluster), but with little overlap between the strains targeted by each. A third group is nested within cluster II according to our analysis, and contained killing activities that were relatively more selective (average of ~7). The fourth group clusters closely with the third group mentioned above due to the relatively low number of targets, but contains strains that have a minimal number (one or two) of targets. The fifth cluster is composed of strains with no observed killing activity.

**Figure 2.**
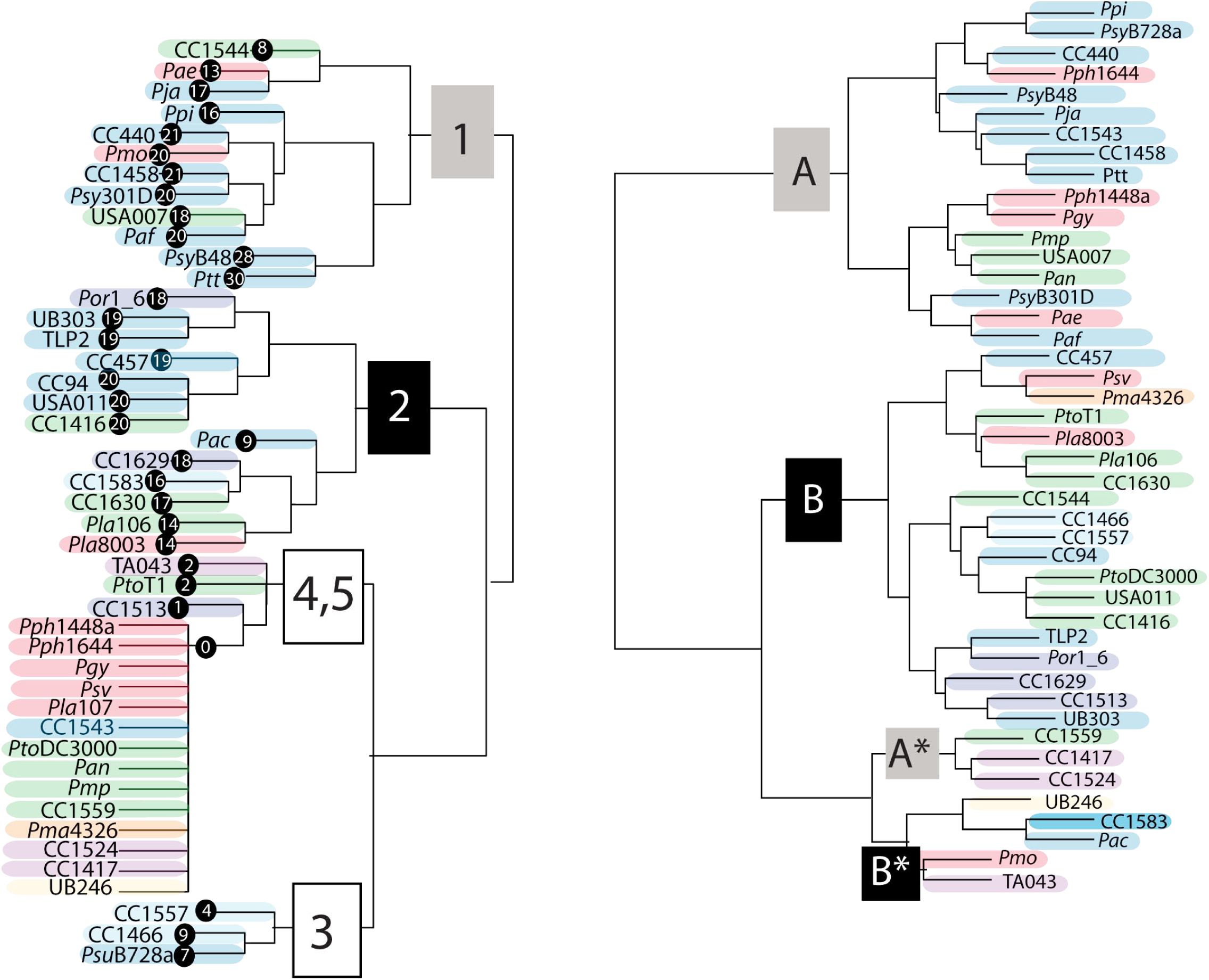
R-type syringacin killing and sensitivity frequently switch across strains. Strains were hierarchically clustered using the phenotypes of killing activity (left) and sensitivity (right) to PEG selected supernatants. Strains are color coded by phylogroup as per Figure 1. Killing spectra clusters (1,2,3,4,5) are shown at nodes that differentiate these groups, and the number of other strains killed by each particular strain (without counting *Pla107* due to inconsistent activity) is shown to the right of the strain name. On the right, strains are clustered by sensitivity to killing, with clusters denoted with letters (A,B,A*,B*).

It is also apparent that, even though many strains have broadly similar killing spectra, there are numerous instances where comparison of these spectra within clusters and groups reveals subtle differences (Figures 1 and 2). For instance, PsyB48 and pathovar *aptata (Ptt)* have the broadest killing activities of any of the sampled strains, targeting 28 and 30 other strains respectively. These strains can both kill the same 24 strains in our screen, however PsyB48 can kill an additional 4 strains (USA007, *Pja, Pmo, Pmp)* and *Ptt* can kill an additional 6 strains (UB246, Pph 1644, CC440, *Ppi, Paf,* TA043). Likewise, while CC440 and CC1458 can both kill 21 strains, only 19 of these targets are shared by both strains as CC440 differentially targets TA043 and UB246 while CC1458 differentially targets *Pmo* and PsyB301D. The clustering of killing spectra between strains further demonstrates both that closely related strains (according to core genome sequences and phylogroups) can have very different killing spectra and conversely that relatively distantly related strains can have convergently evolved spectra.

### Rapid switches of syringamycin killing spectra across strains

Comparison of the *P. syringae* phylogeny reported in Figure 1 with R-type syringacin killing activity reported in Figure 2 demonstrates that killing spectra evolve quite rapidly across strains. Although phylogenies built from whole genome sequences recapitulate traditionally recognized phylogroup structures (see colored shading for phylogroups in Figure 1), hierarchical clustering of killing spectra demonstrates that even closely related strains have dramatically divergent phage derived syringacin targeting capabilities. Rapid diversification is most apparent by comparing coherent phylogenetic clustering of each phylogroup (represented by single blocks of color throughout the phylogeny) with the same colors represented in in hierarchical clusters of activity. For example, compare strains PsyB301D and USA011, which are relatively closely related according to phylogenies built from housekeeping loci, and which target the same number of strains within our assays, but which have very different killing spectra.

### Diverse patterns of R-type syringacin sensitivity

We have also performed hierarchical clustering on strain sensitivity to PEG precipitated antimicrobial compounds (Figure 2). It is also noteworthy that almost all of the *P. syringae* strains were able to be killed by at least one other strain within our collection, with the exceptions being CC1583 *(Pac* is killed by a diffusible compound, but not an R-type syringacin). While, subjectively, there appears to be slightly more phylogenetic congruity overall when considering sensitivity compared to killing spectra (i.e. phylogroup IV strains cluster together for sensitivity but not for killing activity), it is clear that the results form strong parallels with patterns of killing activity described above. There are two main clusters of sensitivity classes, which we term here A and B. We also note two sub-clusters (A2 and B2), which group separately from the main clusters. We refer to these as subclusters because their sensitivity spectra are subsets of those found within main clusters A and B. It is likely that A2 and B2 group together in this analysis, and not with their related parent clusters, because they each are targeted by a small number of strains and clustering is based on phenotypic distance. Therefore, we believe that A2 and B2 cluster together because their patterns are dominated by the lack of activity compared to strains within A and B, and differences in the small number of activities is not enough to override this signal.

There are multiple cases of closely related strains where sensitivity dramatically differs between closely related strains. For instance, strain CC457 from sensitivity group B is closely related to both *Paf* and *Ptt* (which are both sensitivity group A). Likewise, USA007 is classified in sensitivity group A but is closely related to CC1544 and CC1416 from sensitivity group B. There are also numerous cases where relatively divergent strains have convergently evolved to have very similar sensitivities to syringacins. Only a small subset of strains screened herein appear to have the exact same sensitivity spectrum, and there are multiple instances where there are a small number of differences between strains with similar sensitivity spectrums.

### Localized recombination is apparent for a subset of R-type syringacin structural genes

Genome sequences are available for each of the strains screened as part of this study (see <u>10.6084/m9.figshare.5738523</u>), and it is straightforward to comb through this genomic information in order to identify genotypic changes strongly correlated with switches in killing activity. Given that receptor binding proteins (Rbp) have been implicated in targeting of R-type pyocins, analysis of their sequences is a clear first step towards identifying determinants of R-type syringacin specificity. As a test of whether diversity in Rbps correlates with killing activity, we inferred phylogenetic relationships for these predicted receptor binding loci and compared these trees to those inferred for multiple loci found in close genomic proximity and surrounding the receptor binding proteins in the genome *(trpC/trpD* as well as a predicted phage tail fiber protein and tail sheath). As highlighted by colors representative of phylogenetic placement of strains into phylogroups in Figures 1 and 2, phylogenies built from sequences of loci bracketing the receptor binding protein largely recapitulate that inferred from housekeeping gene sequences (Figure 3A). Although the positions of a handful of nodes change, overall strains form coherent groups that match phylogroup assignments. Phylogenetic patterns found on either side of the Rbp are therefore strongly consistent with vertical inheritance of the R-type syringacin locus across the diversity of *P. syringae.* Strikingly, phylogenies built from the receptor binding protein sequences display extensive differences from those inferred from sequences of conserved housekeeping genes, as well as those inferred from sequences for other phage derived syringacin structural genes. For the receptor binding locus, coloring of strain names and thus phylogroup placement of strains is not monophyletic and is instead quite mixed. Moreover, phylogenies built from the sequences of receptor binding proteins form two clear clades and strain membership in these clades is consistent with groupings of killing activity shown in Figure 2.

**Figure 3.**
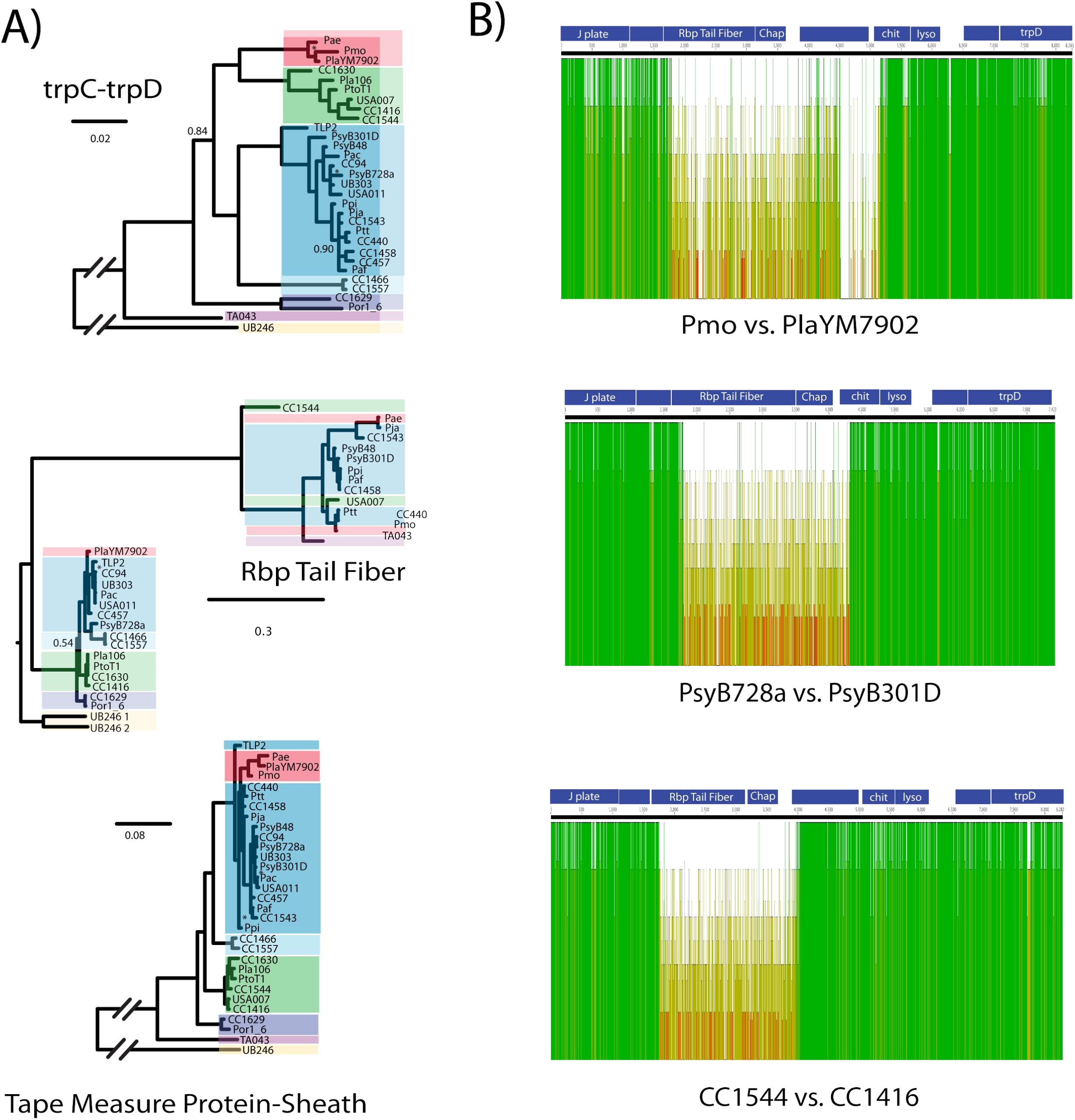
Localized recombination drives diversification of R-type syringacin killing activity. A) Phylogenies are shown for genes found “upstream” (TrpC/TrpD) and “downstream” (Tape measure protein-Tail sheath) of the receptor binding protein for all strains that showed killing activity consistent with R-type syringacins without our assays (with the exception of CC1583, for which the Rbp sequence is truncated). A phylogeny is also shown for the Receptor binding protein itself. Strain names are color coded by phylogroups as shown in figure 1. Posterior probabilities <0.95 are noted with either the actual number or an “*” where space was limited. B) Shown are nucleotide sequence alignments for regions of the genome including the receptor binding protein and chaperone pairs, as well as surrounding syringacin structural genes and *trpD.* Predicted reading frames for genes found in this region are highlighted in blue at the top of each alignment. Green bars indicate high similarity in nucleotide sequence, whereas yellow bars indicate lower similarity. In each case, we find low nucleotide similarity for the area including most of the receptor binding protein and chaperone and much higher similarity for regions upstream and downstream of these two genes.

Differences in phylogenetic placement for the receptor binding protein is clearly suggestive of extensive localized recombination involving receptor binding proteins compared to other surrounding loci that code for structural loci for R-type syringacins. To better gauge the breakpoints and extent of these recombination events, we mapped nucleotide diversity for genomic regions involving these genes from three pairs of strains that are closely related but which cluster into different groups based on killing spectra (Figure 3B; CC1544 and CC1416 from phylogroup I, PsyB728a and PsyB301D from phylogroup II, and *Pmo* and Pla7092 from phylogroup III). Conservation of nucleotides is shown by green blocks within Figure 3B, while divergence is shown by yellow blocks. Surprisingly, the N-terminus of the Rbp is relatively highly conserved across strains but levels of sequence conservation repeatedly shift from high to low at a point inside the predicted Rbp open reading frame. This N-terminal sequence therefore likely acts as one anchor point for recombination events that diversify R-type syringacin killing activity. On the other side, there is no consistent pattern for conservation of nucleotide diversity downstream of the Rbp and chaperone, which suggests that the other recombination breakpoints likely occur in different positions in each of these pairs. However, in each case, it appears as though the breakpoint is upstream of the J-plate protein (a structural protein in R-type syringacin production). Therefore, even though it is difficult to identify the ancestral state for any of these sequences, in each of the three independent pairwise comparisons the receptor binding protein and chaperone have been cleanly switched through recombination and this switch is correlated with a complete overhaul of the killing spectra.

### Transfer of two genes is sufficient to change phage derived syringacin killing spectrum

To test that recombination of the Rbp and chaperone alone were sufficient to retarget syringacins, we utilized a mutant strain of *P. syringae* pv. *syringae* B728a (PsyB728a) in which both the Rbp and chaperone were previously deleted (Hockett et al., 2015). We have already demonstrated that we can complement phage derived syringacin production and killing in this strain background by ectopically expressing the PsyB728a genes from a plasmid (Hockett et al., 2015). However, instead of complementing this strain with its native receptor binding protein and chaperone, we instead chose to test whether killing activity of this strain could be complemented by both loci from a strain with a distinctly different killing spectrum *(P. syringae* pv. *japonica, Pja).* As shown in Figure 4, not only can the receptor binding protein and chaperone from *Pja* complement deletion of these genes from PsyB728a, but the killing spectrum of this complemented strain is consistent with the spectrum from *Pja* instead of *Psy*B728a. Therefore, horizontal transfer of both the receptor binding protein and chaperone is sufficient to retarget *P. syringae* syringacin

**Figure 4.**
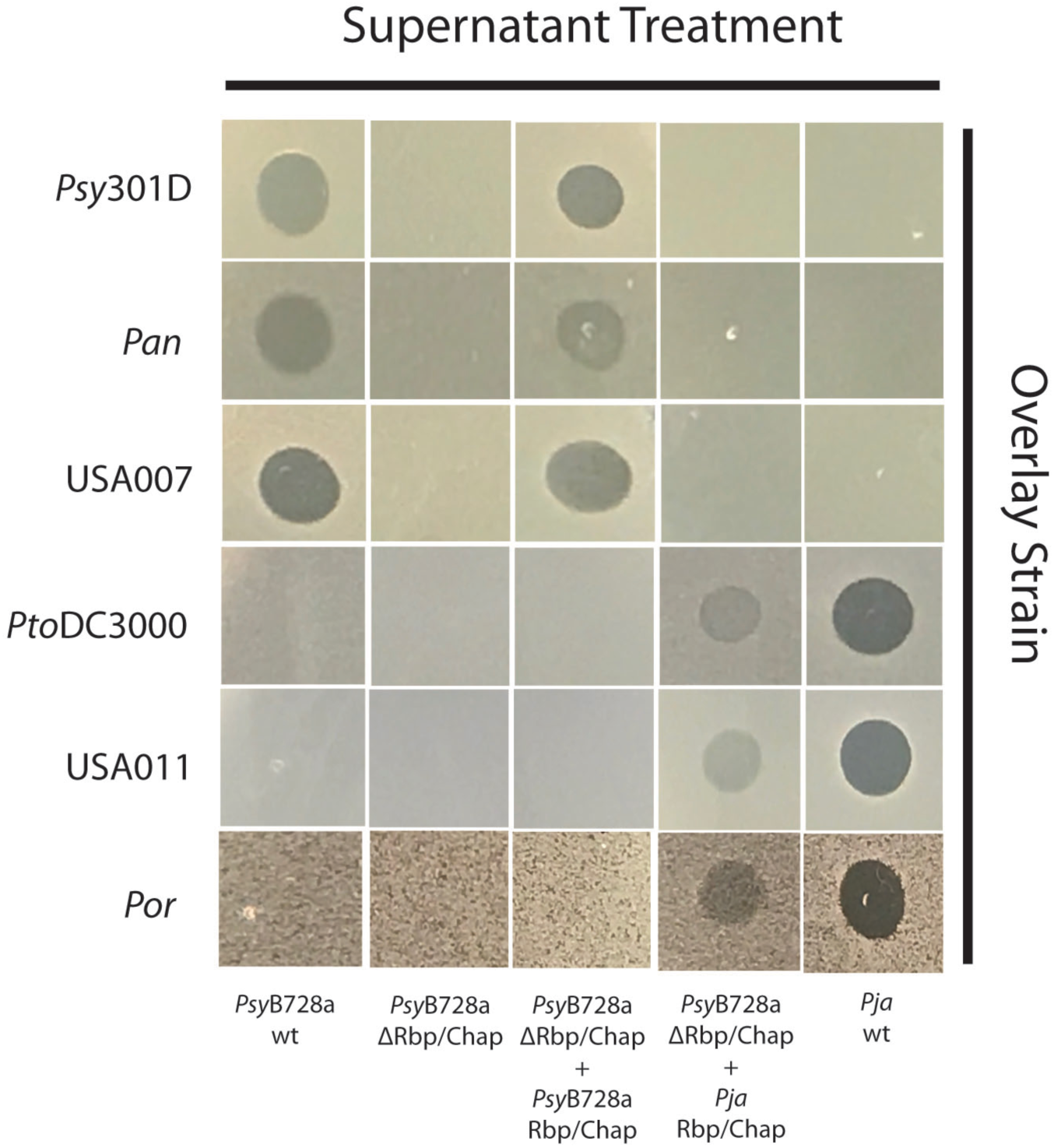
Ectopic expression of the Rbp and chaperone is sufficient to retarget R-type syringacins. The results of overlays of six different strains and five different PEG selected supernatants are shown. Overlay strains are shown on the Y-axis, whereas supernatants are shown on the X-axis. Three strains are sensitive to killing by R-type syringacins produced by PsyB728a and not *Pja* (Psy301D, *Pan,* USA007), while three strains are sensitive to R-type syringacins produced by *Pja* and not PsyB728a (PtoDC3000, USA011, Por1_6). Killing activity is abolished in a mutant of PsyB728a where both the receptor binding protein (Rbp) and chaperone from the R-type syringacin has been deleted, and this mutant can be complemented by expression of both these genes ectopically from a plasmid. Ectopic expression of the Rbp and chaperone from *Pja* in this PsyB728a ARbp/chaperone background retargets this strain so that the killing spectrum matches that of *Pja*.

## Discussion

Phage derived bacteriocins possess great potential as antimicrobials and may play significant roles in structuring bacterial communities under natural conditions (Ghequire and De Mot, 2015; Hawlena et al., 2012; Riley and Wertz, 2002). Given the potential for strong selective pressures to diversify killing spectra, and likewise evolutionary pressure to gain resistance, bacteriocins are also likely to be hotspots for coevolutionary interactions between closely related strains. In order to shed light on evolutionary histories of phage derived bacteriocins in the phytopathogen *P. syringae* (termed R-type syringacins), we performed a screen to measure the spectra of killing activity across a diverse collection of strains and have used genome sequence information to match phenotypic changes to underlying genetic differences.

### Phage derived bacteriocins are the main intra-species killing activity for *P. syringae*

Our results demonstrate that R-type syringacins are the dominant inducible bacteriocin produced across these strains, but also highlight rapid evolutionary shifts in both syringacin host range as well as resistance. Broadly, we find that both R-type syringacin killing spectra and sensitivity to R-type syringacin killing across many strains can be grouped into main two clusters for the strains assayed (1 and 2 for killing, A and B for sensitivity). Moreover, membership in each cluster is highly correlated: Killing spectra cluster I is typically found within strains that are members of sensitivity cluster B and targets stains from cluster A, while killing spectra cluster 2 is typically found within strains that are members of sensitivity cluster A and target strains from cluster B. There are two strains that don’t fit cleanly into killing groups 1 or 2, but instead reflect a mix of both of these (PsyB48 and *Ptt).* Aside from strains that could not easily be grouped into killing and sensitivity clusters, there are three strains that seem to be exceptions to these correlations *(Pmo,* CC1544, PsyB728a). Notably, each of these strains is resistant to its own R-type syringacin even though they are sensitive to other tailocins from the same killing group. Our groupings differ from the previous groupings reported by Vidaver and Buckner (Vidaver and Buckner, 1978), and there are several reasons to think our groupings would differ from those previously reported. First, we clustered strains with similar, but not necessarily identical, killing spectra into the same killing group using a metric of phenotypic distance, whereas in the previous work subtle differences in killing spectra resulted in distinct bacteriocin producer groupings. Second, our assays focused largely on R-type syringacin-mediated killing, while the previous work did not distinguish between R-type and S-type syringacin activity. Finally, the previous work focused largely on a collection of strains belonging to pathovar *syringae* (phylogroup II). Thus, their collection could have potentially teased apart more nuanced interaction by deeper sampling within a narrower phylogenetic range of strains.

### R-type syringacin islands are largely syntenic and vertically inherited across *P. syringae*

We have inferred phylogenies for multiple genes found throughout the R-type syringacin locus for the strains screened as part of this study. Phylogenies for numerous structural genes involved in syringacin production match that inferred from whole genome sequences (at least at the level of phylogroups), which suggests those overall the genomic island for phage derived syringacin production resides in between *trpD* and *trpE* and has been vertically inherited throughout most of the strains classified as *P. syringae* (Figure 3A). The one clear exception to this result, as demonstrated both by phylogenetic divergence as well as genomic context of the syringacin island itself, is phylogroup XIII strain UB246. Structural genes for production of R-type syringacins for this strain are exceptionally divergent from those in other *P. syringae* strains, much more so than the rest of the genome (Figure 2). Furthermore, assembly of this strain’s genome sequence demonstrates that the R-type syringacin locus is not found in between *trpD* and *trpE,* but instead is present at a different (and currently unidentified) genomic location. Thus, it appears as though this strain either maintains a syringacin locus (or has independently acquired this locus) independently from other *P. syringae* strains. Perhaps not coincidentally, this strain also maintains a divergent type III secretion system than other *P. syringae* strains also at a different genomic location than the canonical *P. syringae* locus (Baltrus et al., 2017). It will be interesting for future studies to tease out whether these parallel genomic alterations are merely coincidental, or are the product of reacquisition of both syringacin and a type III secretion system after adaptation to a new niche for this phylogroup XIII strain. Regardless, these patterns suggest that the ability to inject effector proteins and the ability to produce phage derived syringacins goes hand in hand, and potentially speaks to correlated yet currently unrefined ecological role for both of these pathways under natural conditions.

### Independent absence of R-type syringacin killing activity across *P. syringae*

There are numerous instances where surveyed *P. syringae* strains have no R-type syringacin activity across this screened collection. Further investigation suggests that, at least in a subset of cases, that losses of activity have independently occurred and are the product of different underlying genetic causes. Two strains from phylogroup IX show no activity consistent with R-type syringacins, and further inspection of their genome sequences suggests that they lack all of the genes known to be associated with R-type syringacins of *P. syringae*. Indeed, there is only one gene predicted to be found between *trpD* and *trpE* for these strains and there are no other regions of these draft genomes that appear to endode phage derived bacteriocins. This genomic context matches that found in strain UB246, which, combined with their phylogenetic placement (Figure 1), suggests that this may be the ancestral context for this region in *P. syringae*. Therefore, the ability to produce phage derived bacteriocins from a locus in between *trpD/trpE* has either been lost or has never been acquired by this group. Coupled with diversity in the locus from phylogroup XIII strain UB246 noted above, these patterns suggest interesting evolutionary and genomic dynamics for strains considered to be outgroups to those strains traditionally classified as *P. syringae* sensu stricto.

Conversely, although many closely related strains that belong to P. syringae phylogroups I ( 4 of 10 strains) and III (4 of 7 strains) do not appear to produce R-type syringacins targeting strains screened for sensitivity in this study, these strains do possess some structural genes for R-type syringacin production. These strains have at least a semblance of the genetic potential to create phage derived bacteriocins, but either lack the ability to make a subset of required proteins or target a limited number of strains that aren’t represented in our screen. In some cases, we have previously noted instances where genomes appear to lack crucial proteins for the production of these molecules and it is likely that these truly are nulls (Hockett et al., 2015). However, we also note that genome sequences for strains suggest that there are mutations that truncate critically important proteins (i.e. in *Psv,* the chaperone protein has an early stop codon (data not shown)).

### Localized recombination involving the receptor binding protein and chaperone drive changes in R-type syringacin killing spectra

Even though R-type syringacins are composed of multiple proteins, we find that diversity in targeting capability can at least partially be explained by changes in a pair of genes that encode a receptor binding protein and its chaperone, the former being critical for target cell recognition. Moreover, we have demonstrated that acquisition of just these two genes can lead to changes in targeting capability for syringacins for at least one pair of strains. While recombination is known to occur within *P. syringae,* recombination events are usually dominated by events between strains within the same phylogroup (Dillion et al., 2017; McCann et al., 2017). Given sampling biases inherent in these data, it’s difficult to make quantitative statements on rates of recombination. However, comparison of phylogenies within Figure 3, and specifically the multiple cases where strains from different phylogroups share closely related Rbp alleles, suggest that recombination events can occur broadly across phylogroups. Moreover, further comparison suggests that recombination events involving these loci can also occur within phylogroups (i.e. Rbp of Psy301D). Phylogenetic patterns for the Rbp genes suggest that recombination events occur in the least at the level of single open reading frames across the strains sampled here, but it remains a possibility that recombination has occurred over smaller regions within these bacteriocin loci. Essentially, the presence of such recombination events would imply that smaller pieces of the Rbp/chaperone pairs could be swapped in a modular way rather than whole loci. Inspection of sequence data within strains screened here suggests that intra-locus recombination can take place, although our current sampling scheme lacks power and levels of within phylogroup sampling for evidence of such events to emerge. Taken together, patterns in both the frequency and the nature of recombination strongly support the idea that evolutionary dynamics of the Rbp and chaperone loci differ from much of the rest of the core genome.

*P. syringae* genomes are highly plastic in terms of gene presence and absence, but acquisition of loci through horizontal gene transfer is thought to occur mainly through phage transduction or conjugation (Baltrus et al., 2017; Dillion et al., 2017). It is difficult to imagine how such high levels of site specific recombination can be achieved through either conjugation or transduction across these strains though, although It is possible that ssDNA containing these loci could be packaged and transferred along with other phage (Lee et al., 1999). To this point, production of ssDNA has been witnessed from R-type pyocin loci but the ability of such particles to be packaged and transferred remain unexplored. Localized recombination with clean replacement of extant alleles, as seen with the receptor binding protein and chaperone loci, is more likely to occur through natural transformation of extracellular DNA followed by homologous recombination rather than site specific recombination. While there is one report of natural transformation in *P. syringae* occurring *in planta (Lovell et al., 2009),* strains have typically been recalcitrant to recombination of extracellular DNA under laboratory conditions and the signals that trigger competence for natural transformation remain undefined. Transformation may also be mediated through the production of vesicles (Renelli et al., 2004). At present given the weight of all data, although these recombination patterns are easily explained by natural transformation of *P. syringae,* exact mechanisms enabling such localized recombination across *P. syringae* strains remain unclear.

### Overall patterns in *P. syringae* killing activity

The strains screened as part of this study are fairly representative of diversity for *P. syringae* as a whole, and thus we believe that this study provides an adequate view into overall trends for *P. syringae* in terms of killing spectra and sensitivity. Although our strains were biased towards the well studied phylogroups I, II, and III, our screen included representatives from 9 of the 13 acknowledged phylogroups (Berge et al., 2014). One major trend that stands out is that phylogroup II strains appear to possess an above average tendency to be able to kill other strains, while phylogroup I and III strains contain an overabundance of “non-producers”. While it remains a possibility that this pattern is due solely to sampling bias, and given the caveat that “non-producers” may just have a more limited target range, we have included a minimum of 8 strains from each phylogroup reflecting overall evolutionary diversity and phytopathogenic host range of these groups. It may be a coincidence, but strains from phylogroup II also tend to be isolated more frequently from a variety of environmental sources and are also pathogenic on a more diverse suite of hosts than other phylogroups (Berge et al., 2014). It will be interesting for future studies to address whether this pattern of broader R-type syringacin killing capability is related to host generality and environmental hardiness, or how such capabilities are reflective of ecological niches. Previous studies with R-type pyocins and other bacteriocins have strongly suggested that these molecules are widely used to outcompete other strains under natural conditions (Hert et al., 2005; Lavermicocca et al., 2002). With this idea in mind, the complete lack of R-type syringacins for phylogroup IX strains and to a more limited extent the lack of R-type syringacin activity from other strains also potentially reflects fundamental differences in ecological strategies employed by these isolates compared to other *P. syringae* strains.

Combining phylogenetic analyses with phenotypic assays, at present we believe that our data best reflects that *P. syringae* strains can be split into at least two main activity clusters and at least five coherent groups based on R-type syringacin activity. These clusters are qualitatively defined by a combination of syringacin killing spectra, but also reflect phylogenetic relationships of the underlying syringacin receptor binding proteins themselves. These clusters are dominated by two broad classes of phage derived syringacin targeting genes that are phylogenetically distinct, but which each target a relatively high number of our screened strains in patterns that are largely non-overlapping. A third class of syringacin targeting genes maintain a relatively limited and specific killing spectrum across the strains analyzed (~7 on average). Lastly, the “non-producer” strains is likely a group composed of strains with specific killing activities (but for which we haven’t included potential targets in our screen) as well as strains which are truly “non-producers”. This group could be split based on future studies, either by finding strains for which they can target, or by genomic comparisons of syringacin structural genes combined with moving the receptor binding proteins to known producer strains to show that activity exists if the rest of the syringacin structural proteins are made.

### Subtle differences within killing spectra and sensitivity classes

R-type syringacin killing spectra display qualitative differences across strains. However, within these broadly defined classes, it is apparent that many strains have similar, yet non-identical killing spectra and thus subtly differ in their overall pattern. These phenotypic differences may reflect amino acid differences between receptor binding protein-chaperone pairs across strains, which could render R-type syringacins ineffective against a subset of hosts. It is also possible that there are genes outside of those known to function in targeting, where sequence diversity could alter the ability to attach to target cells or may otherwise affect the efficiency of killing. If this is the case, receptor binding protein-chaperone pairs with identical sequences may differ in targeting spectra depending on the strain genomic background. To this point, we highlight strains CC1466 and CC1557 which have identical protein sequences for both their receptor binding proteins and chaperones but which differ in killing spectra (Figures 1 and 3A).

In terms of sensitivity, there are multiple strains that have convergently evolved to be sensitive to the same suite of syringacins. We have recently demonstrated that the main target activity of syringacins is the Lipopolysaccharide (LPS), much like R-type pyocins (Hockett et al., 2017; Köhler et al., 2010). Numerous previous studies have investigated LPS diversity across *P. syringae,* as this structure is the target for many antibodies used to serotype strains before the advent of widespread sequence based identification schemes (Ovod et al., 1997; Zdorovenko and Zdorovenko, 2010). These serotyping studies suggested that LPS structures are quite diverse and changeable, and that strains within the species could be split into at least 15 different groups (Ovod et al., 1997). If LPS is the target of these R-type syringacins, it is thus likely that single types of phage derived bacteriocins can target multiple types of LPS found within *P. syringae.* Alternatively, it is possible that LPS structures have converged across strains in specific ways that enable similar sensitivities despite overall variability. This prediction arises from our observations that an R-type syringacin produced by one strain is often able to differentially target strains within the same pathovar but can also kill strains from different phylogroups and pathovars. However, previous serotyping studies often showed that pathovars were often consistently grouped into the same serotype by LPS pattern and that different pathovars were often grouped into different serotypes (Ovod et al., 1997). From an evolutionary perspective, convergence or diversification of LPS structures could be important when considering trade-offs to syringacin resistance, because the LPS is known to be recognized as a MAMP by plant immune responses and thus changes in the LPS could directly affect virulence *in planta (Ranf et al., 2015).* Likewise, inherent LPS structures likely constrain which types of syringacin loci are able to be acquired by strains because self-targeting could occur if the “wrong” alleles of receptor binding proteins and chaperones are introduced to strains through recombination. The dangers of similar evolutionary tradeoffs were clearly demonstrated in a strain of *P. aeruginosa* evolved to form better biofilms, which changed LPS and left strains open to self-killing by pyocins (Penterman et al., 2014).

### Caveats

We have presented a large-scale screen of general bacteriocin activity across diverse *P. syringae* strains, followed by selection of R-type syringacin activity from these same samples. Although we are confident that much of the activity is as reported, there remain a couple of caveats for these results. First, we note that lysogenic bacteriophage will likely be induced by the same mitomycin C treatments as bacteriocins (Lamont et al., 1989). While we already discounted phage activity wherever single plaques were apparent during the overlay assays, It is possible that some of the reported activity is due to phage that are at high enough concentrations that single plaques blend together and therefore converge upon bacteriocin killing phenotypes. It is also possible that PEG selection may not dilute these phage particles to levels to allow visible plaques. If we have scored phage activity as bacteriocin activity, this will lead to an overestimation of bacteriocin killing, however we remain confident given overall patterns that a far majority of the activity reported is due to bacteriocins.

Weak killing activity in any of the assays is challenging to score, and it is possible that very weak activity (due to a low number of killing particles or due to overgrowth by the overlay strain) could be scored as a null in our assays. To counter this, we have repeated all inductions, PEG selections, and overlays at least twice independently and have reported results where only 1 of 2 assays showed activity. We also note that as with any selection, recovery of killing particles after PEG treatment is not 100% efficient and dilution of phage derived bacteriocins will likely occur. It is therefore possible that lack of killing activity in after PEG treatment could be attributed to R-type syringacins that have simply been diluted to the point where killing in overlay assays is difficult to identify. In particular, the killing spectra of strain CC1543 stands out because there is clearly activity in supernatants and none seen after PEG selection. Furthermore, the actual killing spectra seen in supernatants of CC1543 is close to that of R-type syringacin cluster 1, even though this activity didn’t precipitate with PEG. Moreover, the predicted Rbp sequence from CC1543 is also most closely related to other alleles from strains classified in cluster 1. There don’t appear to be many other instances where killing activity could be improperly scored as that of diffusible bacteriocins, we note that some instances where spectra are subtly different may be due to very weak interactions scored as nulls.

## Conclusions

We have demonstrated that phage derived bacteriocin killing activities can be grouped into multiple clusters, and yet within each cluster there exists subtle differences from strain to strain in terms of activity. Furthermore, sensitivity to phage derived bacteriocins across strains can also be grouped into multiple clusters, the composition of which is highly correlated with each strain’s killing activity. We have shown that killing spectra for phage derived bacteriocins can change rapidly between closely related strains, and that localized recombination of just two genes appears to be sufficient to explain some transitions between killing groups. Even though recombination is known to occur within *P. syringae,* typically it is thought to occur between strains within phylogroups. However, some of the recombination events witnessed here inherently must have occurred between recipient and donor strains from different phylogroups. Switches between killing and sensitivity also evolve frequently between strains, and could speak to larger selective pressures under natural conditions.

It is possible that the interactions reported in this manuscript are highly relevant to understanding structure within natural communities of *P. syringae.* Given the strong selective pressures of bacteriocin killing, it will be interesting to investigate how both killing and sensitivity differ throughout space and time and across different communities on different host plants. In the very least, our data provide a blueprint for evaluating expectations for intermicrobial interactions driven by phage derived bacteriocins within species.

## Materials and Methods

### Strains and Growth Conditions

Genome sequences for all wild type *Pseudomonas syringae* strains used in this study are publically available, and assemblies can be found are listed in <u>10.6084/m9.figshare.5738523</u>. Strains and plasmids constructed for experiments performed in this study are listed in Table 1. For basic maintenance and propagation, all strains were grown on King’s B media (KB) unsupplemented with antibiotics and incubated at 27°C. Where applicable, antibiotic were used at the following concentrations: tetracycline 10 ug/mL, rifampicin 50 ug/mL, nitrofurantoin 40 ug/mL.

**Table 1.** Strains and Plasmids Constructed for this Study.

| **Strain** | **Description** | **Reference** |
| --- | --- | --- |
| *Psy* B278a | *P. syringae* pv. *syringae* B278a |  |
| *Pja* | *P. syringae* pv. *japonica* MAFF301072 |  |
| *Psy*B728a ΔRrbp/KLH5 | *Psy* B728a with clean deletion of Rbp and chaperone (*psyr*_4584 and *psyr*_4585) | Hockett et al. 2015 |
| *Psy*B728a ΔRrbp+pDAB296 / KLH24 | *Psy*B728a ΔRrbp complemented with *Psy*B728a Rbp and chaperone | Hockett et al. 2015 |
| *Psy*B728a ΔRrbp+pRbp-B728a / KLH27 | *Psy*B728a ΔRrbp complemented with *Psy*B728a Rbp and chaperone | Hockett et al. 2015 |
| *Psy*B728a ΔRrbp+pRbp-*Pja* / KLH41 | *Psy*B728a ΔRrbp complemented with *Pja* Rbp and chaperone | This paper |
| **Plasmid** | **Description** | **Reference** |
| pBV226 | Gateway destination vector with expression driven by *nptII*, TetR | Vinatzer et al. 2006 |
| pDAB296 | MTN1970-2 EV construct in pBV226, TetR | Baltrus et al. 2012 |
| pRbp-B728a | Rbp and chaperone ORFs from *Psy*B728a in pBV226, TetR | Hockett et al. 2015 |
| pRbp-*Pja* | Rbp and chaperone ORFs from *Pja* in pBV226, TetR | This paper |

### Bacteriocin Induction and Preparation

The basic protocol for overlay assays has been described previously (Hockett and Baltrus, 2017). Briefly, for bacteriocin production, a single colony of the strain of interest is picked to 2–3mL KB and grown overnight in a shaking incubator at 220rpm and 27°C. The next day, strains were back diluted 1:100 in 3mL of KB media and grown for 3–4 hours. At this point, mitomycin C was added at 0.5 μg/ml concentration to induce bacteriocin production and cultures were incubated with shaking overnight. After this point, cultures were centrifuged at 20,000 × g for five minutes, supernatant was taken and added to a new tube. To sterilize the supernatant chloroform was added at 100μl per 1mL, vortexed for 15 seconds and incubated at room temperature for 1 hour. The supernatant was then centrifuged at 20,000 × g for five minutes. The upper aqueous layer was removed without any of the bottom chloroform layer, and then top layer was incubated in the fume hood for a minimum of four hours to allow residual chloroform to evaporate.

### Polyethlyne Glycol (PEG) Precipitation

Separation of high molecular weight R-type syringacins from the lower molecular weight bacteriocins compounds was performed by PEG precipitation. This was done by adding NaCl and PEG 8000 to final concentrations of 1M and 10% (w/v), respectively. The supernatants were then left overnight at 4°C or incubated for 1 hour on ice. Centrifugation then took place for 30 minutes at 4°C. The supernatant was removed and the pellet was resuspended in 100–250 ul of buffer (10mM Tris, 10 mM MgSO4, pH 7.0). Residual PEG was removed by two sequential equal volume extractions with chloroform. Residual chloroform was allowed to evaporate from uncapped samples for four hours in a fume hood.

### Soft Agar Overlay Protocol

A single colony of the strain being used as an overlay was picked to 2–3mL KB and grown overnight in a shaking incubator at 220rpm and 27^o^C. The next day, strains were back diluted 1:100 in 3mL of KB media and grown for 3–4 hours. The 0.4% agar was melted and 3ml was added to a culture tube, once it had cooled down 100ul of the strain of interest was added. It was then poured over a petri dish containing kb media, and swirled to cover the entire dish. It was allowed to solidify for 15–20 minutes and then the 2ul of supernatant and PEG prepped samples were spotted onto the dish. They were allowed to grow 1–2 days and then the presence and absence of killing activity was recorded. All assays were performed twice, with two independent bacteriocin inductions (and subsequent PEG selection) and overlays per supernatant-overlay combination. A majority of the activity was repeatable across assays, but assays were repeated for a third time if results were inconsistent. If no clear result emerged after three assays (i.e. strong, weak, and no killing across these three assays), data was coded as presence of “killing” but with a qualifier notation. Full killing data on interactions from across assays can be found here: <u>10.6084/m9.figshare.5734125</u>

### Hierarchical Clustering

A smaller killing-sensitivity matrix was created using only the data from PEG prepped supernatants, with presence of killing activity coded as a “1” and absence of activity coded as a “0”. In cases where activity was inconsistent across assays, data was coded as a “1”. All data on killing of *Pla107* was excluded from this cluster analysis because of variability caused by a secondary killing activity due to megaplasmid present within this strain. The PEG-only killing matrix, *sans* Pla107, can be found here:<u>10.6084/m9.figshare.5734122</u>. Calculation of phenotypic distances between killing profiles for each strain was performed in R (version 3.3, (R Core Team, 2014)) using the “dist” function and “euclidean” method. Hierarchical clustering was then performed on phenotypic distances ugn the “hclust” function and “ward.D2” method. R scripts used for these clustering calculations can be found here: <u>10.6084/m9.figshare.5738427</u> Profiles for sensitivity were calculated in the same way, except that data in the original PEG-only killing matrix was transposed. This transposed matrix can be found here: <u>10.6084/m9.figshare.5734119</u>

### Phylogenetics

Nucleotide and protein sequences of all genes involved were identified through tblastn searches of available genomes. For MLSA phylogenies in figure 1, nucleotide sequences for fragments of *gyrB, rpod, gltA, gapA,* and *acnB* were concatenated and then aligned. For phylogenies in figure 3A, amino acid sequences for fragments of TrpC/TrpD and tail sheath/tape measure protein were independently concatenated and aligned. Amino acid sequences of receptor binding proteins were aligned separately from the sequences. We have excluded the amino acid sequence of CC1583 from this analysis because it was truncated within the draft genome assembly and we could not identify the C terminus for this allele. In all cases, sequences were aligned using the ClustalX (Thompson et al., 2002) with default parameters, with alignments polished by hand if necessary. Alignments for all sequences used to infer phylogenies can be found at <u>10.6084/m9.figshare.5734128</u>, <u>10.6084/m9.figshare.5734131</u>, and <u>10.6084/m9.figshare.5734134</u>.

All phylogenies were inferred using MrBayes 3.0 (Ronquist and Huelsenbeck, 2003). For MLSA phylogenies, convergence occurred after 1,000,000 generations and a burn in of 250,000 was used. For TrpC/TrpD, convergence occurred after 1,000,000 generations and a burn in of 250.000 was used. For tail sheath/tape measure protein phylogenies, convergence occurred after 1.500.000 generations and a burn in of 375,000 was used. For Rbp phylogenies, convergence occurred after 2,500,000 generations and a burn in of 625,000 was used.

### Retargeting of R-type Syringacin Killing Activity

A construct (start codon to stop codon, see sequence at <u>10.6084/m9.figshare.5738436</u>) containing the receptor binding protein and its chaperone from *P. syringae* pv. *japonica* MAFF301072 was amplified from genomic DNA of this strain and recombined into pDONR207 using BP clonase (Invitrogen, Carlsbad CA) to create plasmid pKLHECO24. pKLHECO24 was purified and the Rbp/chaperone construct was recombined into pBAV226 using LR recombinase (Invitrogen, Carlsbad CA) to create pKLHECO26. The final plasmid, in which the *Pja* Rbp and chaperone are cotranscribed from an *nptII* promoter, was mated into KLH5 (ΔRrbp from Hockett et al. 2015), and a tetracycline resistant colony was isolated. Overlays were performed as above, with all strains grown in unsupplemented KB media prior to induction. Whole pictures, cropped to create images for Figure 4, are available here: <u>10.6084/m9.figshare.5734137</u>.

